# The genomic and ecological context of hybridization affect the probability that symmetrical incompatibilities drive hybrid speciation

**DOI:** 10.1101/233924

**Authors:** Aaron A. Comeault

**Affiliations:** Biology Department, University of North Carolina, 250 Bell Tower Road, Chapel Hill, 27599, USA.

**Keywords:** hybridization, hybrid speciation, reproductive isolation, epistasis

## Abstract

Despite examples of homoploid hybrid species, theoretical work describing when, where, and how we expect homoploid hybrid speciation to occur remains relatively rare. Here I explore the probability of homoploid hybrid speciation due to “symmetrical incompatibilities” under different selective and genetic scenarios. Through simulation, I test how genetic architecture and selection acting on traits that do not themselves generate incompatibilities interact to affect the probability that hybrids evolve symmetrical incompatibilities with their parent species. Unsurprisingly, selection against admixture at ‘adaptive’ loci that are linked to loci that generate incompatibilities tends to reduce the probability of evolving symmetrical incompatibilities. By contrast, selection that favors admixed genotypes at adaptive loci can promote the evolution of symmetrical incompatibilities. The magnitude of these outcomes is affected by the strength of selection, aspects of genetic architecture such as linkage relationships and the linear arrangement of loci along a chromosome, and the amount of hybridization following the formation of a hybrid zone. These results highlight how understanding the nature of selection, aspects of the genetics of traits affecting fitness, and the strength of reproductive isolation between hybridizing taxa can all be used to inform when we expect to observe homoploid hybrid speciation due to symmetrical incompatibilities.

## Introduction

Modern genomic data and analyses are revealing that naturally occurring hybridization and admixture between divergent lineages is not rare (Maqbool *et al*. 2015; Racimo *et al*. 2015; Pease *et al*. 2016; Wallbank *et al*. 2016). The evolutionary consequences of hybridization are however diverse. On one hand, hybridization has been described as “the grossest blunder in sexual preference which we can conceive of an animal making” (Fisher 1930). On the other, hybridization can be a generative force, facilitating adaptive evolution via adaptive introgression (Song *et al*. 2011; Dasmahapatra *et al*. 2012) or promoting diversification through hybrid speciation (Anderson & Stebbins 1954; Buerkle *et al*. 2000; Gross & Rieseberg 2005; Mallet 2007). Cases of hybrid speciation exist (Rieseberg *et al*. 2003; Gompert *et al*. 2006; Duenez-Guzman *et al*. 2009; Salazar *et al*. 2010; Nice *et al*. 2013; Hermansen *et al*. 2014; Lamichhaney *et al*. 2017), and some have suggested that hybridization may be responsible for a larger fraction of species diversity than previously appreciated (Mallet 2007; Mavarez & Linares 2008). However, linking the specific mechanism(s) through which hybridization causally leads to the evolution of reproductive isolation (RI) between hybrids and their parents, in many putative cases, remains a major challenge (Schumer *et al*. 2014).

Hybrid speciation can occur either with or without a change in ploidy between hybrid lineages and their parents (Stebbins 1959; Rieseberg *et al*. 1995; Hegarty & Hiscock 2005; Mallet 2007). Polyploid hybrid speciation is rare in animals, but relatively common in plants (see Stebbins 1959; Hegarty & Hiscock 2005), because, relative to plants, incidence of polyploidy are rare in most groups of animals (Orr 1990; Otto & Whitton 2000; Mable 2004). By contrast, homoploid hybrid speciation (HHS) has been shown to occur in plants (e.g. *Helianthus anomalus;* (Rieseberg *et al*. 1995, 2003; Ungerer *et al*. 1998), animals (e.g. *Heliconius heurippa;* (Jiggins *et al*. 2008; Melo *et al*. 2009; Salazar *et al*. 2010), and fungi (Leducq *et al*. 2016). Additional examples of putative homoploid hybrid species are becoming more common (reviewed in Gross & Rieseberg 2005; Mavarez & Linares 2008). For details of specific examples of hybrid species, I refer the reader to citations presented throughout this manuscript; hereafter I focus specifically on the processes generating RI during HHS.

At least four studies have quantitatively explored conditions that can lead to HHS. These studies demonstrate that admixed populations are more likely to stabilize, and evolve RI from their parental species, when they display a high rate of selfing (in plants; McCarthy et al. 1995) or assortative mating (in animals; Duenez-Guzman et al. 2009), show transgressive segregation at traits influencing fitness in a novel environment (Buerkle *et al*. 2000), and/or are geographically isolated from their parental species (McCarthy *et al*. 1995; Buerkle *et al*. 2000; Schumer *et al*. 2015). Each of these factors can promote reproductive isolation between admixed and parental lineages and allow for genomic stabilization and independent evolution to occur within admixed populations. In addition to cohesion through geographic, ecological, or sexual isolation, hybrid populations can display intrinsic incompatibilities with their parental species (Rieseberg *et al*. 1995; Hermansen *et al*. 2014). These intrinsic incompatibilities can help maintain stable hybrid populations despite the opportunity for ongoing gene flow with their parental species. In order to better appreciate when hybridization is most likely to drive speciation, it is therefore important to understand the conditions and mechanisms that result in genomic stabilization within hybrid lineages, and the evolution of RI between hybrid lineages and their parents.

One such mechanism is when two or more independently acting genetic incompatibilities fix for alternate parental genotypes in a hybrid population. This ‘balancing’ of incompatibilities results in admixed genomes (or more specifically, haplotypes) that are compatible with each other, but will manifest at least one incompatibility with either of their parental species (herein referred to as “symmetrical incompatibilities”). Loci that can generate symmetrical incompatibilities include chromosomal rearrangements (McCarthy *et al*. 1995; Buerkle *et al*. 2000) or epistatic pairs of loci that affect fitness as a result of inter-allelic interactions (e.g. Dobzhansky-Muller Incompatibilities) (Schumer *et al*. 2015). For example, consider a pair of loci that interact through epistasis and are segregating for both parental ancestries at equal frequencies. Under the assumptions that selection favors interactions between alleles sharing the same ancestry within each pair symmetrically (e.g. Table 2) and that the strength of selection is greater than drift (i.e. greater than ~1/(2*N_e_*)), both parental ancestries have an equal probability of fixing within each of the two pairs of interacting loci. Extending this example to multiple independent pairs of ‘epistatic loci’, the probability of fixing for either parent 1 or parent 2 alleles across all epistatic pairs is 2×0.5^*n*^, where *n* is the number of epistatic pairs. Conversely, the probability of evolving mixed ancestry and some amount of RI due to symmetrical incompatibilities across the *n* epistatic pairs is 1 − (2×0.5^*n*^). All-else being equal (e.g. independent assortment of loci and no selection acting on additional traits), symmetrical incompatibilities may therefore readily evolve in sufficiently admixed populations (Schumer *et al*. 2015).

McCarthy et al. (1995) and Buerkle et al. (2000) tested the probability that symmetrical incompatibilities would evolve between admixed populations and their parents as a result of novel “chromosomally balanced” genotypes with respect to two rearrangements that differed between the parental species. Their simulations show that admixed populations can evolve RI under this mechanism, and that the probability of evolving RI increases both as hybrid fitness in a novel environment and geographic isolation from parental populations increases. Taken with the results presented by Schumer et al. (2015), these analyses describe (1) how symmetrical incompatibilities can evolve in admixed populations and generate RI between admixed and parental populations and (2) suggest that the probability of evolving symmetrical incompatibilities is contingent upon the nature of selection acting on hybrid individuals.

In nature, the fitness of naturally occurring hybrids in different environments relative to their parents is seldom known; however, it is likely to vary depending on multiple factors. In some cases, such as in *Helianthus* sunflowers, hybrids may be more fit than their parental species in certain environments (Rieseberg *et al*. 1995, 2003). In others, hybrids may be less fit than their parents, and this may (or may not) depend on the environment that a hybrid finds itself in (Vamosi & Schluter 1999; Linn *et al*. 2004; Bridle *et al*. 2006; Delmore & Irwin 2014; Turissini *et al*. 2017). It is therefore likely that the evolution of symmetrical incompatibilities will be affected by the specific fitness function acting on admixed genotypes. By extension, selection acting at linked sites will also affect the probability of evolving symmetrical incompatibilities. Understanding the genetic architecture of traits, and the form of selection acting on those traits, is therefore important to fully appreciate the scenarios that either permit or constrain the evolution of symmetrical incompatibilities in admixed populations.

In this manuscript I use forward-time individual-based simulations to illustrate how the nature of selection acting on, and the linkage relationships between, loci that generate incompatibilities (hereafter “epistatic” loci) and those that affect an additional trait under selection (hereafter “adaptive” loci) affect the probability that admixed populations evolve symmetrical incompatibilities. To accomplish this, I simulate three different types of selection acting on adaptive loci and varied (1) the strength of selection acting on both adaptive and epistatic loci, (2) the order of loci along a chromosome, and (3) recombination rates between adjacent loci. Each of these parameters were varied in a ‘hybridizing deme’ experiencing gene flow from demes containing their parental species. Consistent with previous work, these simulations show how selection favoring admixed genotypes at adaptive loci tends to increase the probability of evolving symmetrical incompatibilities, while selection favoring alleles from one or both parental species at adaptive loci tends to decrease the probability of evolving symmetrical incompatibilities. Both the strength of selection acting on the different types of loci and their genetic architecture affect the probability that a hybrid population will evolve symmetrical incompatibilities. Below I summarize these effects and highlight how understanding how selection acts on hybrids, along with knowledge of the genetic basis of traits that are subject to selection and underlie reproductive isolation between parental species, can be used to predict when we expect to observe homoploid hybrid species evolve.

## Materials and Methods

### General Description of Model

I carried out forward-time simulations of demes composed of 1,000, diploid individuals. Hybrid populations in nature seldom evolve without some level of ongoing hybridization with parental populations; therefore, I simulated structured populations that consisted of two ‘parental demes’ and a central ‘hybrid deme’. Hybridization occurred in the hybrid deme that experienced immigrants from the two parental demes at rate *m*, per parental deme. I simulated three different rates of *m*: 0.0001, 0.001, and 0.1, corresponding to an average of 0.1, 1, and 10 immigrant individuals from each parental deme per generation, respectively. Simulations were initiated under each of two different conditions: (1) the hybrid deme was composed of equal proportions of randomly mating parental genotypes or (2) the hybrid deme was composed of an equal number of males and females that were heterozygous with respect to ancestry across all loci (i.e. all individuals were F_1_ hybrids).

Each individual’s genome consisted of a single chromosome with seven equally-spaced loci (Figure 1). Two pairs of loci were subject to selection due to epistasis. (Two is the minimum number of pairs required to allow for symmetrical incompatibilities to evolve.) The remaining three loci additively affected an individual’s fitness in the environment (e.g. ecological, social, or sexual environment). The relative fitness of an individual was a function of their genotype at these loci (see “Selection” below; Tables 2 and 3). I tracked allele frequencies at each locus, within each population, for 1,000 generations, recording allele frequencies every 10 generations. Mating was accomplished by randomly sampling individuals, with replacement, with the probability of sampling an individual being proportional to their fitness. All simulations were carried out using Python scripts (available at https://github.com/comeaultresearch/simuHybrid) that utilize objects and functions contained within the simuPOP environment (Peng & Kimmel 2005).

**Figure 1.**
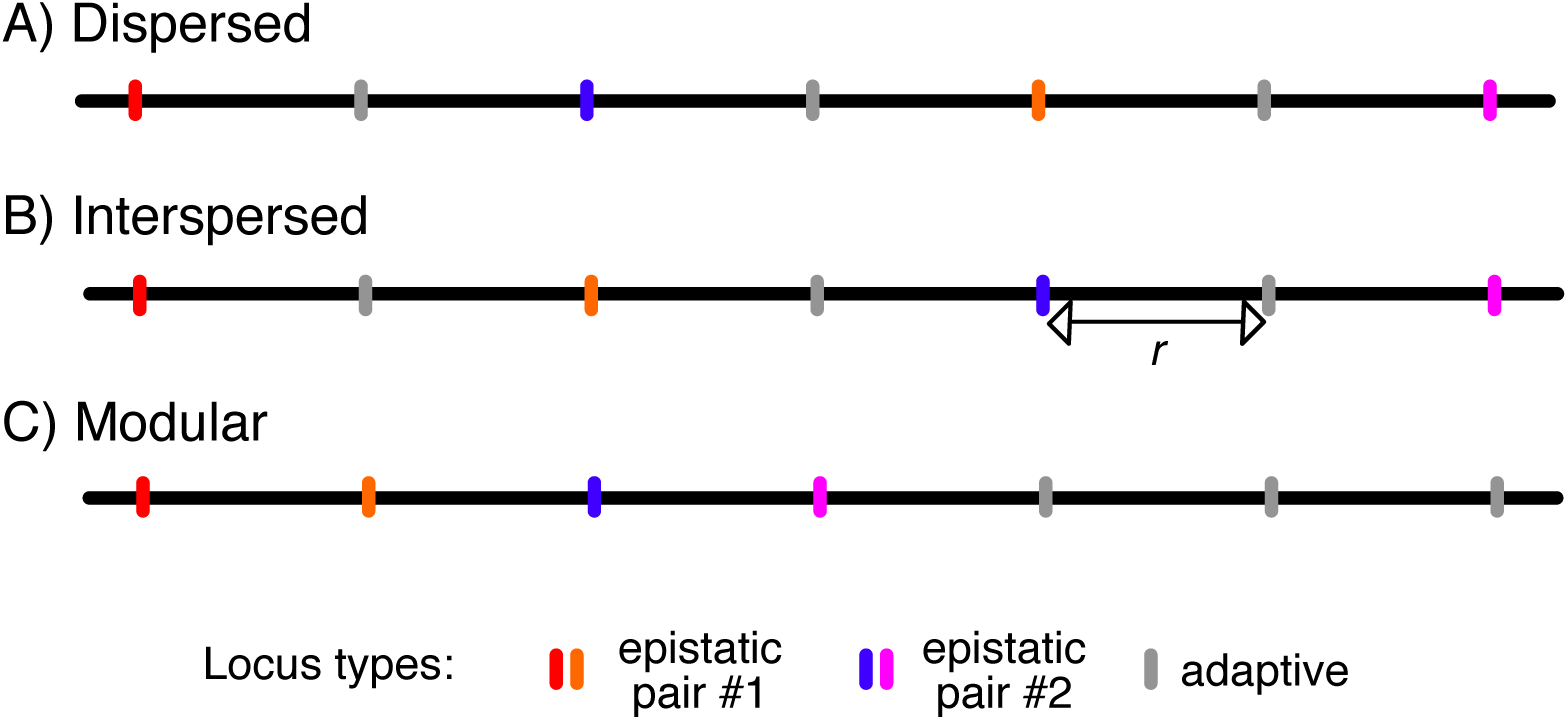
Illustration of the three genetic architectures simulated in this study. Each horizontal black line represents a haploid chromosome and vertical lines indicate the position of loci. Recombination occurs along the chromosome at a rate of *r* between adjacent loci.

### Genetic Architecture

Loci were equally spaced along each individual’s chromosomes. The two pairs of loci that contain epistatically-interacting loci (i.e. “epistatic” loci) affected the fitness of an individual as described in Table 1. The effect that these loci have on fitness is solely due to epistasis. Epistatic loci may represent incompatibilities that, for example, cause sterility, but may also underlie any trait that depends on the interaction between multiple loci to function properly. The three other loci additively affect the fitness of an individual as described in Table 2 (i.e. “adaptive” loci). These loci can be thought of as affecting any trait that is controlled by additively acting genetic effects. Adjacent loci recombined at a rate of 0.1, 0.2, or 0.5 per generation. The recombination rates of 0.1 and 0.2 allowed me to test the effect of linkage on the evolution of symmetrical incompatibilities. The maximum rate of recombination (0.5) allowed for random assortment of loci and is equivalent to each locus being located on its own chromosome.

**Table 1.** List of variables and parameters used for simulating evolution within hybrid swarms.

| Variable / Parameter | Description | Values used |
| --- | --- | --- |
| $N$ | Total number of diploid individuals within each population. | 1,000 |
| $n$-loci | Number of diploid loci: two pairs of epistatic loci; three loci additively affecting fitness. | 7 |
| generations | Number of generations populations were monitored. | 1,000 |
| $s_{epistatic}$ | Selection coefficient acting against mismatched alleles at epistatic loci. | 0.000, 0.001, 0.01, 0.05, 0.10 |
| $s_{adaptive}$ | Selection coefficient acting against parent #2 alleles at loci additively affecting fitness in the environment. | 0.00, 0.01, 0.02, 0.03, 0.04, 0.05, 0.06, 0.07, 0.08, 0.09, 0.10 |
| $r$ | Recombination rate between adjacent loci. | 0.1, 0.2, 0.5 |
| $m$ | Probability of migration into from parental populations into hybrid zone. | 0, 0.0001, 0.001, 0.01 |
| genetic architecture | Arrangement of epistatic and adaptive loci along a chromosome. | 3 different architectures (see Methods and Figure 1). |

**Table 2.** The strength of selection as a function of genotype at a pair of ‘epistatic’ loci. Alleles have ancestry from either parent 1 (*P_1_* alleles) or parent 2 (*P_2_* alleles). Total selection due to maladaptive epistatic interactions (*S_epistatic_*) was summed across the two epistatic pairs considered during simulations. The dominance coefficient (*h*) was held constant at 0.5 in all simulations.

|  |  | genotype at locus 1 |  |  |
| --- | --- | --- | --- | --- |
| | | $P_1P_1$ | $P_1P_2$ | $P_2P_2$ |
| genotype at locus 2 | $P_1P_1$ | 0 | $h^*s_{epistatic}$ | $2^*s_{epistatic}$ |
| | $P_1P_2$ | $h^*s_{epistatic}$ | $2^*h^*s_{epistatic}$ | $h^*s_{epistatic}$ |
| | $P_2P_2$ | $2^*s_{epistatic}$ | $h^*s_{epistatic}$ | 0 |

In addition to varying recombination rates, I tested how the physical arrangement of loci along a chromosome affects the probability of evolving symmetrical incompatibilities. I either positioned loci such that the distance between similar types of loci was maximized (“dispersed” genetic architecture; Figure 1A), the two epistatic pairs were on opposite ends of the chromosome, but were interspersed by the adaptive loci (“interspersed” genetic architecture; Figure 1B), or loci were grouped by type such that epistatic loci and pairs were adjacent to each other and were not interspersed by an adaptive locus (“modular” genetic architecture; Figure 1C).

### Selection

During simulations, an individual produced offspring proportional to their relative fitness. An individual’s fitness was a function of selection acting against alleles subject to either epistatic (*S*_epistatic_) or ‘adaptive’ selection (*S*_adaptive_) such that ω = 1 − (*S*_[epistatic pair 1]_) − (*S*_[epistatic pair 2]_) − (*S*_adaptive_). Selection acted independently on each epistatic pair, and the number of ‘mismatched’ alleles within a given pair determined fitness (Table 2).

I simulated three different models of selection on adaptive loci. First I simulated ‘directional selection’, where selection on the three adaptive loci acted additively and alleles with ancestry from one of the parents (hereafter referred to as “P1”) were always favored over alleles with ancestry from the other parent (hereafter “P2”), except in the case where there was no selection acting on these loci (Table 3A). My rationale for simulating this scenario is to expand on treatments of hybrid speciation where hybrids are afforded a fitness advantage in a certain environment (Buerkle *et al*. 2000) or where their fitness is independent of the environment (Schumer *et al*. 2015). The particular parent that I deem selectively favored is arbitrary and represents a scenario where ancestry from one parental species at adaptive loci is favored over the second, while hybrids have intermediate fitness. Second, I simulated ‘diversifying selection’, where selection acted such that homozygous parental genotypes across all three adaptive loci were favored over heterozygous or admixed parental genotypes (Table 3B). This scenario reflects one where hybrid genotypes are at a fitness disadvantage relative to parental genotypes, and parental genotypes are equally fit. Third, I simulated ‘selection-for-admixture’, where selection favored admixed genotypes across the three adaptive loci over parental and heterozygous genotypes (Table 3C). This scenario represents one where hybrids have a selective advantage, such as simulated by Buerkle *et al*. (2000). The difference between the scenario modeled by Buerkle *et al*. and that presented here is that the ‘ecological’ locus in Buerkle *et al*. (2000) segregated independently of the inversions that caused symmetrical incompatibilities, while in this study I explicitly model different scenarios of linkage between adaptive and epistatic loci.This allows me to compare the probability that selection-for-admixture will promote the evolution of symmetrical incompatibilities under different genetic scenarios.

**Table 3.** Descriptions of the three fitness schemes imposed on ‘adaptive’ loci. The total strength of selection against possible genotypes across the three adaptive loci is shown (*S_adaptive_*) along with a description of the different genotypes. Total *S_adaptive_* was subtracted from 1 when determining the relative fitness of an individual during simulation.

| Total $s_{adaptive}$ | Genotype Description |
| --- | --- |
| **A) ‘directional selection’** |  |
| $n_{ALT}(s_{adaptive})$ | Where $n_{ALT}$ is the number of alleles with ancestry from the ‘unfit’ parent. |
| **B) ‘disruptive selection’** |  |
| $n_{MINOR}(s_{adaptive})$ | Where $n_{MINOR}$ is the number of minor ancestry alleles if the number of minor ancestry alleles is less than 3 or all adaptive loci are heterozygous. |
| $5(s_{adaptive})$ | If two loci are homozygous with different ancestry and the third is heterozygous. |
| **C) ‘selection-for-admixture’** |  |
| $6(s_{adaptive})$ | If homozygous for the same ancestry across all adaptive loci. |
| $5(s_{adaptive})$ | If two loci are homozygous for the same ancestry and the third is heterozygous. |
| $n_{HET}(s_{adaptive})$ | Where $n_{HET}$ is the number of heterozygous loci if $> 1$ locus is heterozygous. |
| $1(s_{adaptive})$ | If two loci are homozygous with different ancestry and the third is heterozygous. |
| $0(s_{adaptive})$ | If two loci are homozygous with ancestry from the same parent and the third is homozygous with ancestry from the other parent. |

For epistatic loci, I simulated selection strengths (*S*_epistatic_) of 0, 0.001, 0.01, 0.05, or 0.1. For adaptive loci, selection (*S*_adaptive_) ranged from 0 to 0.1, in increments of 0.01. The maximum total strength of selection I consider is when *S*_epistatic_ = 0.1 and *S*_adaptive_ = 0.1. At this maximum strength of selection, F_1_ hybrids have a relative fitness of 0.3 under each model of selection. Parental genotypes have respective fitness of 1 (P1) and 0.4 (P2), 1, or 0.4 under the directional selection, diversifying selection, and selection-for-admixture models, respectively. The weakest combination of non-zero selection strengths I consider is *S*_epistatic_ = 0.001 and *S*_adaptive_ = 0.01, corresponding to an F_1_ hybrid fitness of 0.966 under each model of selection. At this minimum strength of selection, parental genotypes have a fitness of 1 and 0.94, 1, or 0.94, under the directional selection, diversifying selection, and selection-for-admixture models, respectively. The models of selection and strengths of selection I simulate were chosen to represent biologically plausible scenarios. For example, hybridizations that produce a large fraction of sterile F_1_ offspring (Coyne & Orr 1989; Coyne *et al*. 2004), to those where hybrids show more subtle deficits in traits that affect their ability to survive or procure resources such as food or mates (Blows & Allan 1998; Bolnick & Lau 2008; Delmore & Irwin 2014; Rennison *et al*. 2015; Turissini *et al*. 2017), to those where admixed genotypes are afforded a fitness advantage over their parental species (Rieseberg *et al*. 2003).

### Gene flow

Migration (*m*) was independent of genotype, and individuals from the parental demes moved into the hybrid deme with probability 0.0001, 0.001, or 0.01, for all combinations of *S*_epistatic_, *S*_adaptive_, *r*, and genetic architecture described in Table 1.

### The effect of initial conditions on the evolution of symmetrical incompatibilities

To test how the amount of hybridization occurring in a hybrid zone affects the probability of evolving symmetrical incompatibilities, I initiated simulations either with a hybrid deme containing an equal number of P1 and P2 individuals that mated at random or a hybrid deme containing all F_1_ hybrids. Under both of these starting conditions, I simulated three rates of migration (*m* = 0.0001, 0.001, and 0.01) for all combinations of *S*_epistatic_, *S*_adaptive_, *r*, and genetic architecture described in Table 1. I quantified the effect that a forced bout of hybridization (i.e. all individuals initiated as F_1_ hybrids) had on the evolution of symmetrical incompatibilities by calculating the proportional change in the number of hybrid populations evolving symmetrical incompatibilities under the ‘all F_1_s’ relative to the ‘randomly mating parents’ starting condition.

### Definition of evolving reproductive isolation

I considered a population of hybrids to have evolved RI from their parental species, due to symmetrical incompatibilities, when the difference in mean allele frequency (AF) at the two epistatic pairs of loci was greater than 0.9. This condition represents a scenario where the population is nearly fixed for alleles coming from one parental species at one epistatic pair (e.g. mean P1 allele frequency > 95%) and nearly fixed for alleles coming from the second parental species at the second epistatic pair (e.g. mean P_2_ allele frequency > 95%). I use 90% AF difference as a threshold defining the evolution of RI because the majority of haplotypes within a population that has a difference in parental allele frequency at the two epistatic pairs > 0.9 will be fertile with other hybrids from that population, but manifest incompatibilities with either parental species (the strength being proportion to *S*_epistatic_).

Hybrid speciation differs from ‘classical’ speciation in that barriers to gene flow do not need to evolve *de novo*, potentially leading to rapid speciation. As such, for each population that showed evidence of evolving RI, I recorded the time it took for allele frequencies at the two epistatic pairs to differ by > 0.9, to the nearest 10 generations.

## Results and Discussion

### Selection on epistatic interactions and the evolution of symmetrical incompatibilities

An important parameter that affects the evolution of symmetrical incompatibilities is the strength of selection acting to maintain functional epistatic interactions within independent epistatic pairs (*S*_epistatic_). When I simulated populations initiated with 1000 randomly mating parental individuals (equal proportions) subject to weak (0.001) or nonexistent (0) *S*_epistatic_, little gene flow from parental populations (*m* = 0.0001), moderate linkage between adjacent loci (*r* = 0.2), and no selection on adaptive loci (i.e. *S*_adaptive_ = 0), a maximum of 3 of 500 simulated populations evolved symmetrical incompatibilities, across all three genetic architectures (blue and black points in Figure 2). This is because populations tended to maintain parental diversity at epistatic loci when *S*_epistatic_ was weak (less-than or equal-to 0.001 for the simulations summarized in this manuscript). More generally, when epistatic interactions are subject to weak selection and symmetrical incompatibilities do evolve, the magnitude of RI will also be weak. For example, the reduction in fitness of an offspring produced by a mating between an individual from an admixed population that evolved symmetrical incompatibilities and either parent species would be 0.1% when *S*_epistatic_ = 0.001. The same scenario for *S*_epistatic_ = 0.05 or *S*_epistatic_ = 0.1 would result in a 5 or 10% decrease in fitness, respectively. Therefore, meaningful RI is unlikely to evolve through symmetrical incompatibilities unless parental species have accumulated genetic differences that result in at least moderately strong incompatibilities.

**Figure 2.**
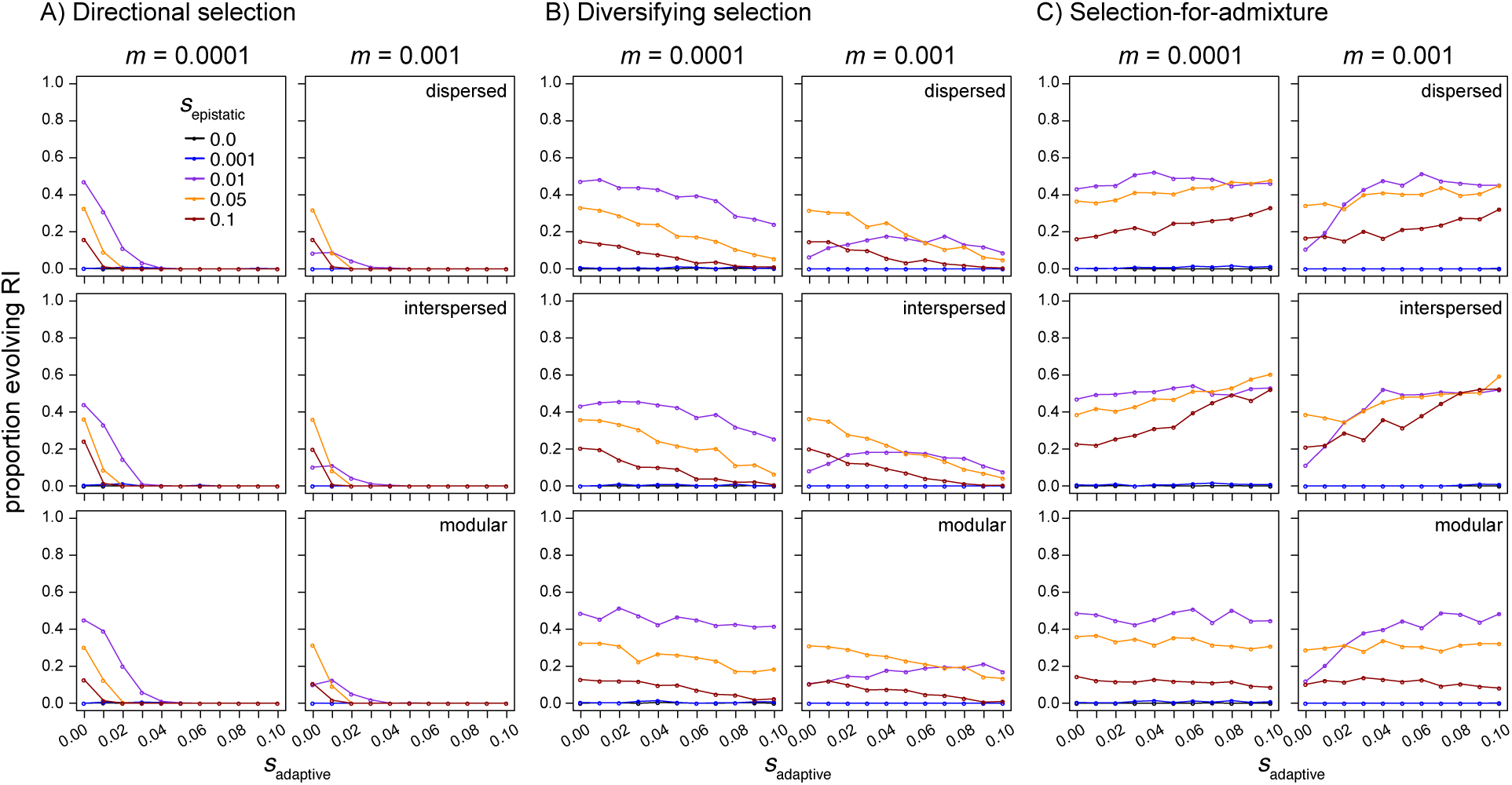
The frequency of hybrid speciation (proportion of 500 simulated hybrid swarms evolving reproductive isolation; y-axis) as a function of the strength of selection acting on epistatic loci (*S*_epistatic_; colored points and lines) and selection acting on additional ‘adaptive’ loci subject to selection (*S*_adaptive_; x-axis; A: directional selection model; B: diversifying selection [i.e. parental genotypes equally favored]; C: selection-for-admixture). Results are shown for hybrid populations simulated with an inter-locus recombination rate of 0.2, migration rates of 0.0001 and 0.001 (panel columns), and with different linear arrangements of loci along the chromosome (i.e. genetic architectures; panel rows).

The strength of *S*_epistatic_ also affects the probability that recombinant haplotypes will persist in a population. When *S*_epistatic_ is strong, recombinant haplotypes are less likely to be maintained in the population and symmetrical incompatibilities are less likely to evolve. For example, when I simulated hybridization in populations experiencing little gene flow from parental populations (*m* = 0.0001) and no selection on additional adaptive loci (*S*_adaptive_ = 0), the greatest proportion of populations evolved RI when sepistatic was moderate (0.01 ; see purple line in left column of panels in Figure 2A-C), with the proportion evolving RI decreasing as the strength of *S*_epistatic_ increased (gold and red lines in left column of panels of Figure 2A-C). This result illustrates how the total strength of selection acting to maintain functional epistatic interactions can reduce the ability of admixed haplotypes to form when species come into secondary contact and hybridize. As such, symmetrical incompatibilities that will contribute to meaningful isolation between admixed and parental lineages are most likely to evolve when *S*_epistatic_ is moderate (relative to *m*; see following section), because weak *S*_epistatic_ will result in variation being maintained within epistatic pairs or generate proportionally weak incompatibilities, while strong *S*_epistatic_ will limit the opportunity for recombinant haplotypes to form.

### Gene flow

As expected, gene flow from parental species generally tends to limit the probability that symmetrical incompatibilities evolve. Specifically, because gene flow can swamp locally adapted epistatic interactions, higher rates of gene flow tend to increase the threshold strength of *S*_epistatic_ required for symmetrical incompatibilities to evolve. For example, consider the purple points between the left and right columns of figure 2A, B, and C: when *S*_epistatic_ = 0.01, fewer populations evolve RI when *m* = 0.001 compared to when *m* = 0.0001. By contrast, for *S*_epistatic_ > 0.01, a similar proportion of populations evolve RI when *m* = 0.0001 or *m* = 0.001 because the relative strength of *S*_epistatic_ is greater than rates of gene flow from parental populations.

Interestingly, with modest gene flow (*m* = 0.001), symmetrical incompatibilities were able to evolve under all three models of *S*_adaptive_ I simulated, as long as selection against hybrids was not too strong (increasing values on the x-axes of Figure 2A and B). This result also depended on the strength of linkage between epistatic and adaptive loci, with tighter linkage further reducing the proportion of populations evolving symmetrical incompatibilities (Figures 4 and S1). By contrast, at high rates of gene flow (*m* = 0.01, or the equivalent of 10 immigrants from each parental population each generation), symmetrically compatibilities were only able to evolve under the directional and diversifying selection models with moderate linkage between loci (*r* = 0.2) when *S*_epistatic_ was strong (0.1; red lines in Figure 3A and B); and even then, the probability they evolved was low (less than 1% of populations). The only exception was that symmetrical incompatibilities evolved with appreciable frequency (> ~20%) in the face of high gene flow when there was selection for admixture and *S*_epistatic_ was strong (i.e. 0.05 or 0.1; gold and red points in Figure 3C). These dynamics illustrate how the probability of evolving symmetrical incompatibilities can remain relatively high (> ~20%), even under high rates of gene flow (i.e. 10 immigrants from both parental species every generation) when selection-for-admixture and *S*_epistatic_ are also strong.

**Figure 3.**
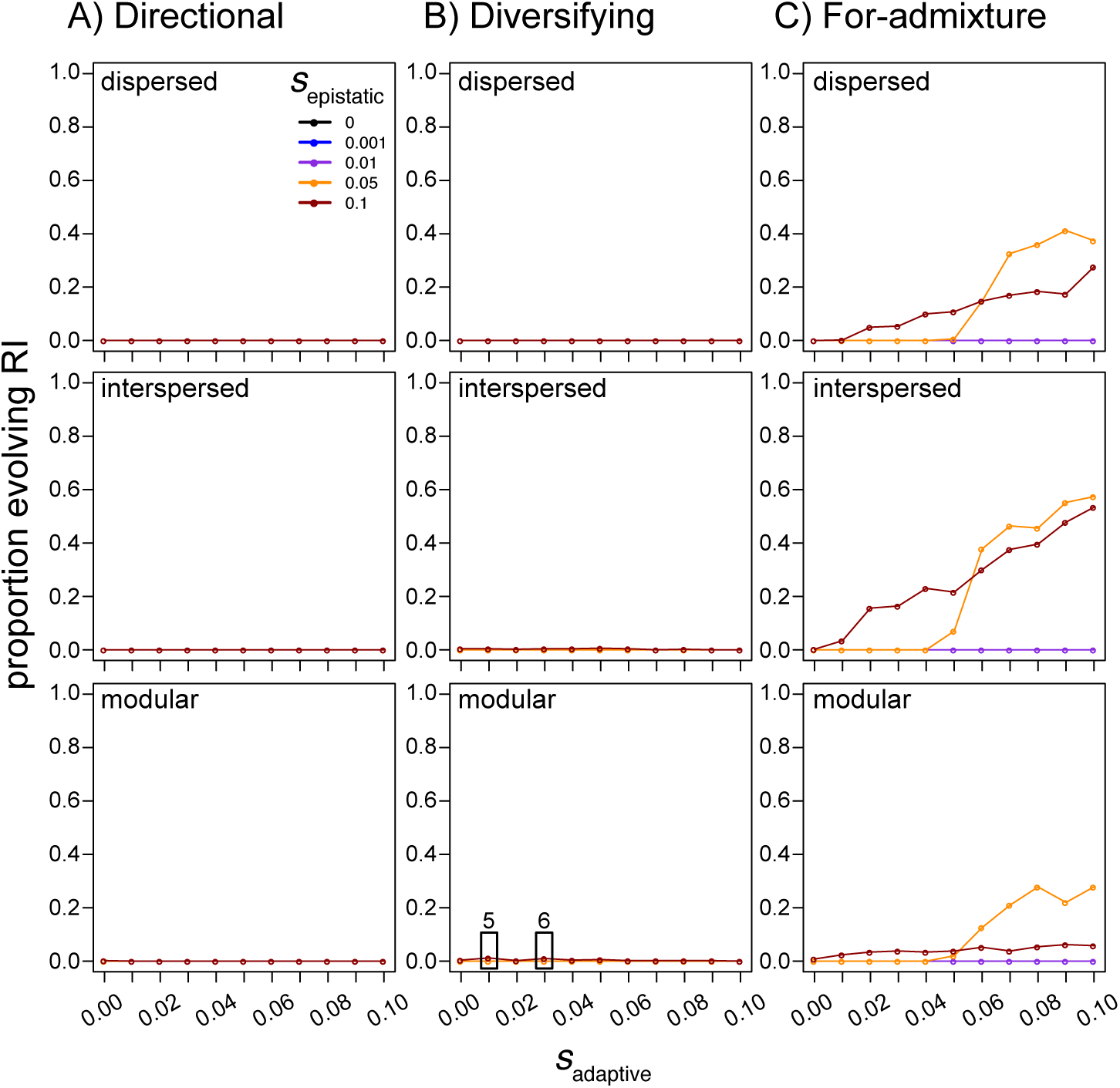
The proportion of hybrid populations evolving symmetrical incompatibilities (y-axis) at high levels of gene flow (*m* = 0.01). Under the directional and diversifying selection models, symmetrical incompatibilities only evolved when *S*_epistatic_ = 0.1, and even then, was rare (less than 1%). Two exceptions are highlighted by black rectangles in the bottom panel of B, with the number of simulated populations that evolved symmetrical incompatibilities given above the rectangles. Panels in C show how symmetrical incompatibilities are most likely to evolve when there is selection-for-admixture and both *S*_adaptive_ and *S*_epistatic_ are strong. Results are shown for hybrid populations simulated with an inter-locus recombination rate of 0.2 and with different linear arrangements of loci along the chromosome (i.e. genetic architectures; panel rows).

**Figure 4.**
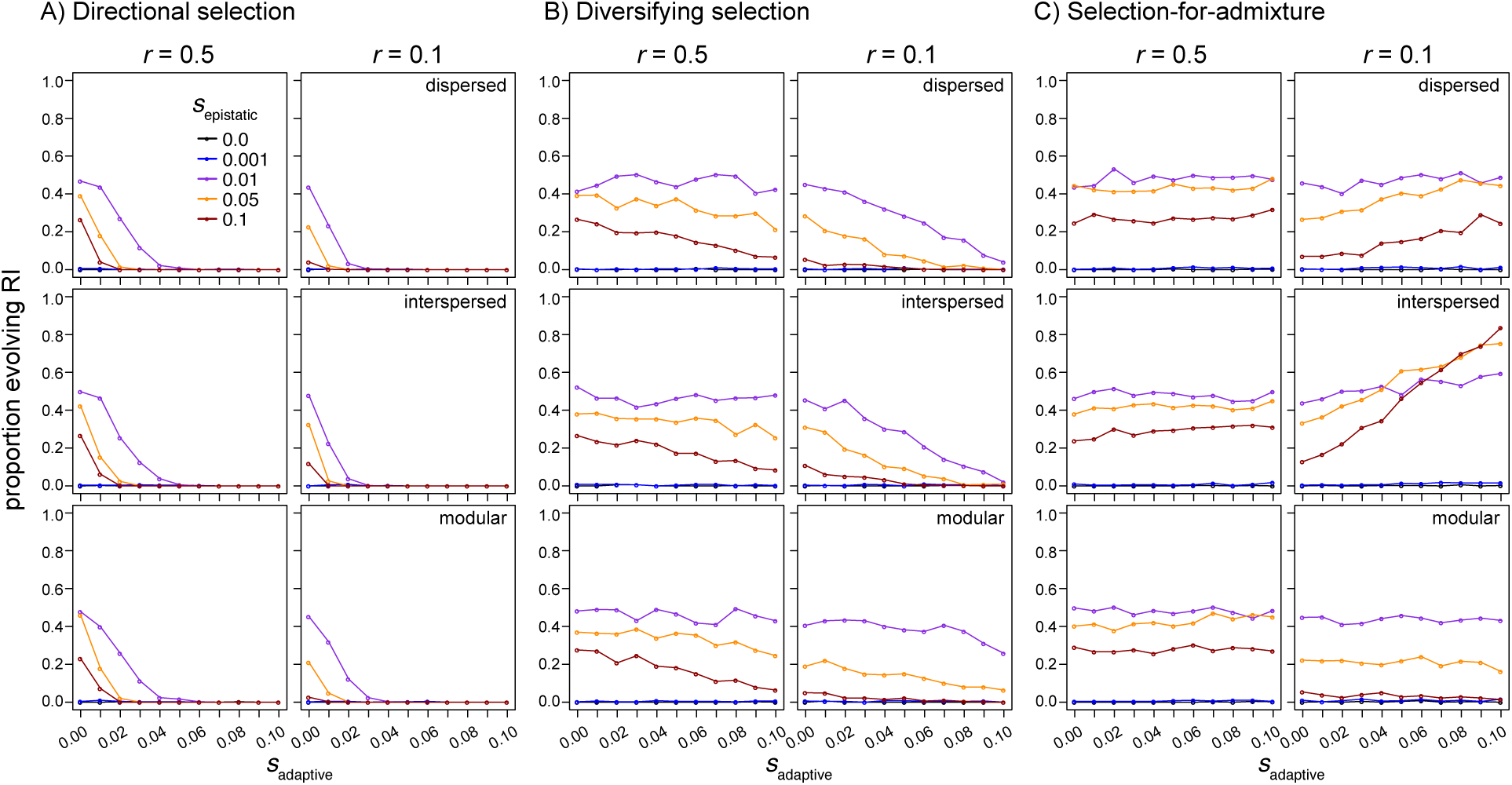
Selection at linked sites and the evolution of symmetrical incompatibilities. Linkage between epistatic and adaptive loci tends to decrease the probability of evolving symmetrical incompatibilities when adaptive loci are subject to directional or diversifying selection (panels in A and B, respectively), but increase the probability of evolving symmetrical incompatibilities when selection favors admixture (C). Results are shown for populations simulated with inter-locus recombination rates of 0.5 (i.e. no linkage; left column of panels) or 0.1 (moderate linkage; right column of panels), *m* = 0.0001, and with different linear arrangements of loci along the chromosome (i.e. genetic architectures; panel rows). Note that genetic architecture is only relevant when *r* is less than 0.5. Refer to Figure S1 for results with *m* = 0.001.

### Selection on adaptive loci and the evolution of symmetrical incompatibilities

In addition to the strength of epistatic selection and rates of gene flow, changes in allele frequencies at epistatic loci can be influenced by selection at linked sites (Maynard Smith & Haigh 1974; Barton 2000). Below I explore the effect of three different models of selection acting on ‘adaptive’ loci linked to the epistatic loci responsible for generating symmetrical incompatibilities. I first present results from simulations initiated with a hybrid deme composed of randomly mating parental species, and then discuss the consequences of a forced bout of admixture in the section *“The effect of initial conditions on the evolution of symmetrical incompatibilities’’*.

Selection acting on sites subject to *S*_adaptive_ either decreased or increased the probability that symmetrical incompatibilities evolved, and the direction of this effect depended on the form of *S*_adaptive_. Directional selection that favored ancestry from one parental species over the other at adaptive loci always reduced the probability that populations of hybrids evolved incompatibilities (Figure 2A). When there is no linkage between epistatic and adaptive loci (*r* = 0.5), this reduction occurs because selection favors ancestry from one parent over the other and limits the opportunity for recombinant haplotypes to form (left column of panels in Figure 4A). Specifically, selection favoring ancestry from one parent over the other at the adaptive loci biases epistatic loci to evolve toward the fitter parent’s ancestry (Figures S2 – S4). This effect was consistent at low, moderate, and high levels of gene flow (Figures 2A and 3A). Under the directional selection model, we therefore expect that as *S*_adaptive_ increases, ancestry within admixed populations will evolve towards the fitter parent and the evolution of symmetrical incompatibilities will be less likely. For the parameter values I simulated, this resulted in no symmetrical incompatibilities evolving when *S*_adaptive_ was greater than 0.03 and there was at least some linkage between adaptive and epistatic loci (Figure 2A and 4A).

When the fitness of parental ancestries is not skewed towards one parent and hybrids are less fit than their parental species (i.e. the diversifying selection model), increasing selection against hybrids (and admixed genotypes) also tends to reduce the probability of evolving symmetrical incompatibilities; however, the magnitude of this effect is much less than for the directional selection model (compare panels between Figure 2A and B). For example, when *S*_adaptive_ is greater than 0.03 and *S*_epistatic_ is greater than 0.001, an appreciable proportion (> 0.1) of admixed populations evolved symmetrical incompatibilities under the diversifying selection model (Figure 2B), while almost none evolved symmetrical incompatibilities under the directional selection model (Figure 2A). Unlike under the directional selection model, the arrangement of loci along the chromosome affected the magnitude of the reduction in the proportion of populations that evolved RI with increasing *S*_adaptive_ under the diversifying selection model (compare down panels in Figure 2A and B). For example, with moderate *S*_epistatic_ (0.01), low migration (*m* = 0.0001), weak linkage (*r* = 0.2), and diversifying selection, as *S*_adaptive_ increases from 0.02 to 0.08 there is a 35%, 30%, and 17% reduction in the proportion of simulated populations that evolve symmetrical incompatibilities for the dispersed, interspersed, or modular genetic architectures, respectively. A modular architecture can therefore facilitate the evolution of symmetrical incompatibilities relative to the dispersed and interspersed architectures when *S*_adaptive_ is strong (yellow and red lines in Figure 2B), migration rates are modest (right panels in Figure 2B), and parents do not differ in their fitness (i.e. under the diversifying selection model).

The two models of selection summarized above both impose selection against hybrid and admixed genotypes at adaptive loci. A third outcome of hybridization is that there is transgressive segregation for fitness-associated traits, resulting in admixed genotypes that are at a selective advantage relative to parental genotypes. Indeed, previous work has shown how symmetrical incompatibilities are more likely to evolve when hybrids have a fitness advantage in a novel environment (see Figure 2 of Buerkle *et al*. 2000), and novel ecological traits in hybrids is a hallmark of one of the best examples of homoploid hybrid speciation: sunflowers in the genus *Helianthus* (Rieseberg *et al*. 1995, 2003). The simulations that I present here recapitulate this result, with the primary difference being that I explicitly simulate linkage between the loci subject to ecological selection (*S*_adaptive_) and those that generate incompatibilities.

Linkage and the ordering of loci along the chromosome (genetic architecture) has the opposite effect on the evolution of symmetrical incompatibilities under the selection-for-admixture model when compared to the directional or diversifying selection models: symmetrical incompatibilities were more likely to evolve under the dispersed and interspersed architectures, on average, than the modular genetic architecture (compare down panels of Figure 2C). (Note that selection-for-admixture only pertains to the adaptive loci and selection acts on epistatic loci the same way in all three models of ‘adaptive selection’.) This result is due to both selection favoring admixed genotypes (in the case where *r* = 0) and linkage between adaptive and epistatic loci in the dispersed and interspersed architectures (when *r* > 0; Figures 2 and 3). Consistent with previous work (Buerkle *et al*. 2000), symmetrical incompatibilities are therefore most likely to evolve when selection favors hybrids, with linkage and genetic architecture interacting to increase the probability that different pairs of epistatic loci evolve to fix different ancestries.

### Time to evolution of RI

Because hybridization requires two species or their gametes to be present in the same location (at least temporarily), the faster that incompatibilities are able to stabilize within admixed populations, the more likely they will show meaningful RI from their parental species in the face of ongoing hybridization. To determine how quickly RI evolved due to symmetrical incompatibilities, I recorded the time (to the nearest 10 generations) it took novel hybrid genotypes to evolve a mean allele frequency difference at the two epistatic pairs of loci greater than 0.9. As expected, the stronger *S*_epistatic_ was, the faster symmetrical incompatibilities tended to evolved (different colored points in Figure 5). Relative to *S*_epistatic_, both *S*_adaptive_ and genetic architecture had negligible effects on the time it took to evolve RI (x-axis of panels and panel columns in Figure 5, respectively). The one exception to this pattern was that increasing *S*_adaptive_ under the selection-for-admixture model resulted in decreasing the time it took to evolve symmetrical incompatibilities when *S*_epistatic_ was moderate (*S*_epistatic_ = 0.01 ; purple points in Figure 5C). This result highlights how once populations begin to evolve allele frequency differences at epistatic pairs of loci, the primary factor affecting the speed that those pairs fix alternate parental alleles is the strength of selection acting to maintain viable epistatic interactions; however, increasing selection on linked loci can increase the speed at which RI evolves in situations where *S*_epistatic_ is not already very strong.

**Figure 5.**
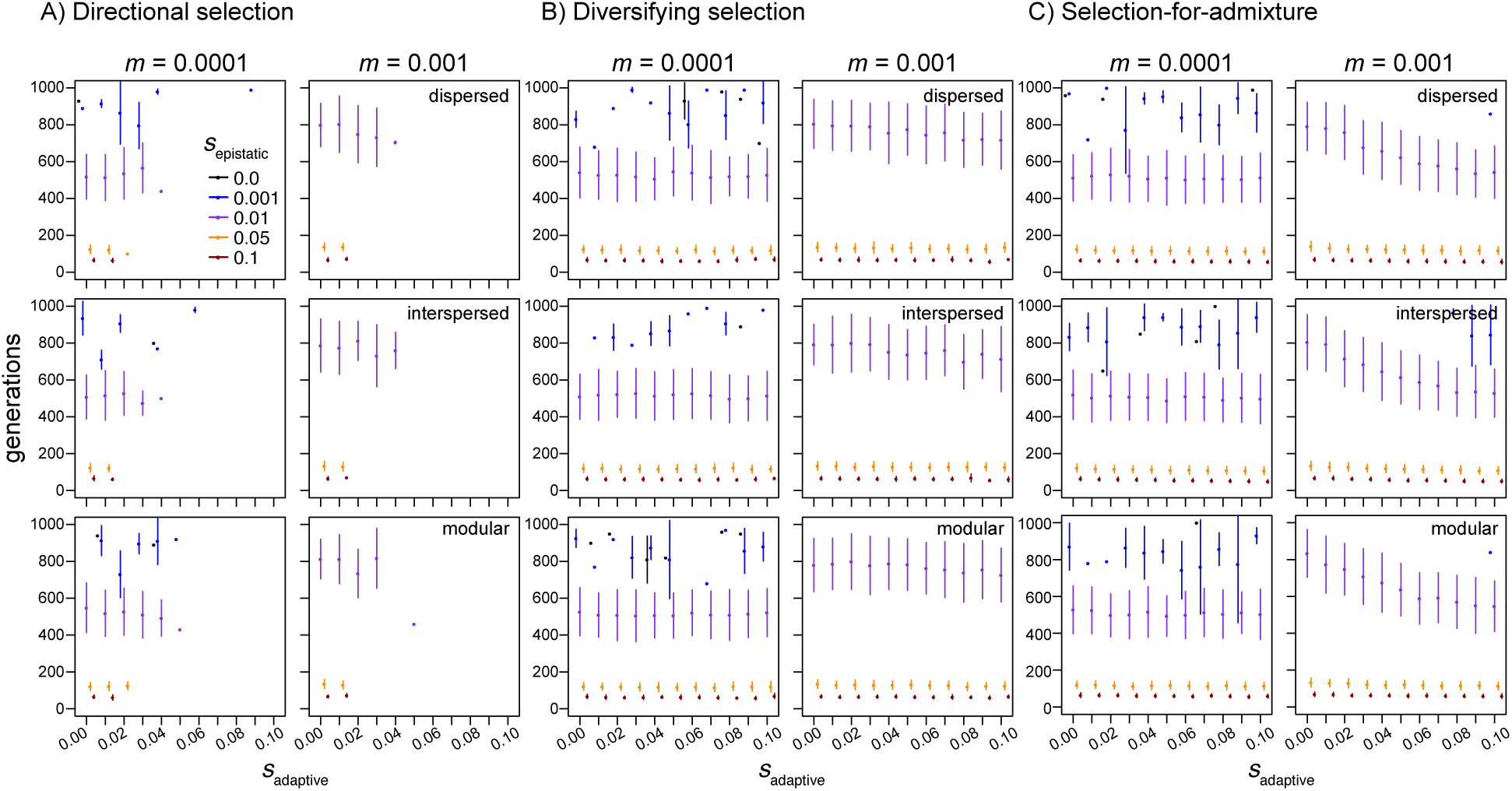
The number of generations required for hybrid populations to evolve reproductive isolation from their parental species. Time is given in generations along the y-axis of each panel for different strengths of selection against alleles at loci affecting fitness in the environment (x-axis). Each colored point within the panels represents the mean time to speciation for hybrid swarms that evolved reproductive isolation from their parental species and points are staggered along the x-axis. Vertical lines are bounded by the 2.5% and 97.5% empirical quantiles of time to speciation for a given set of hybrid populations. Missing points occur for parameter combinations where no populations evolved RI. Results are shown for hybrid populations simulated with an inter-locus recombination rate of 0.2.

### The effect of initial conditionS on the evolution of Symmetrical incompatibilities

When I forced a bout of hybridization by initiating simulations with a hybrid deme composed of F_1_ hybrids, symmetrical incompatibilities were, in general, more likely to evolve than when simulations were initiated with randomly mating individuals of the parental species (Figure 6). This was particularly true when *S*_adaptive_ was greater than zero under the directional or diversifying selection models (Figure 6A and B, respectively). Under directional selection, the relative enrichment in the proportion of populations evolving symmetrical incompatibilities increased as both *S*_adaptive_ and as *S*_epistatic_ increased (compare increasing values on the x-axes and the purple, gold, and red lines in Figure 6A, respectively). By contrast, with selection-for-admixture, an initial bout of hybridization had much less of an effect on increasing the proportion of populations that evolved symmetrical incompatibilities (Figure 6C). In this case, I only observed a modest ~ 1-fold enrichment in the probability of evolving symmetrical incompatibilities when *S*_epistatic_ was very strong (i.e. red lines in Figure 6C). An initial bout of admixture can therefore promote the evolution of symmetrical incompatibilities in scenarios where selection minimizes the probability that recombinant haplotypes will form: i.e. with increasing *S*_adaptive_ and *S*_epistatic_ under the directional or diversifying selection models and with increasing *S*_epistatic_ under the selection-for-admixture model.

**Figure 6.**
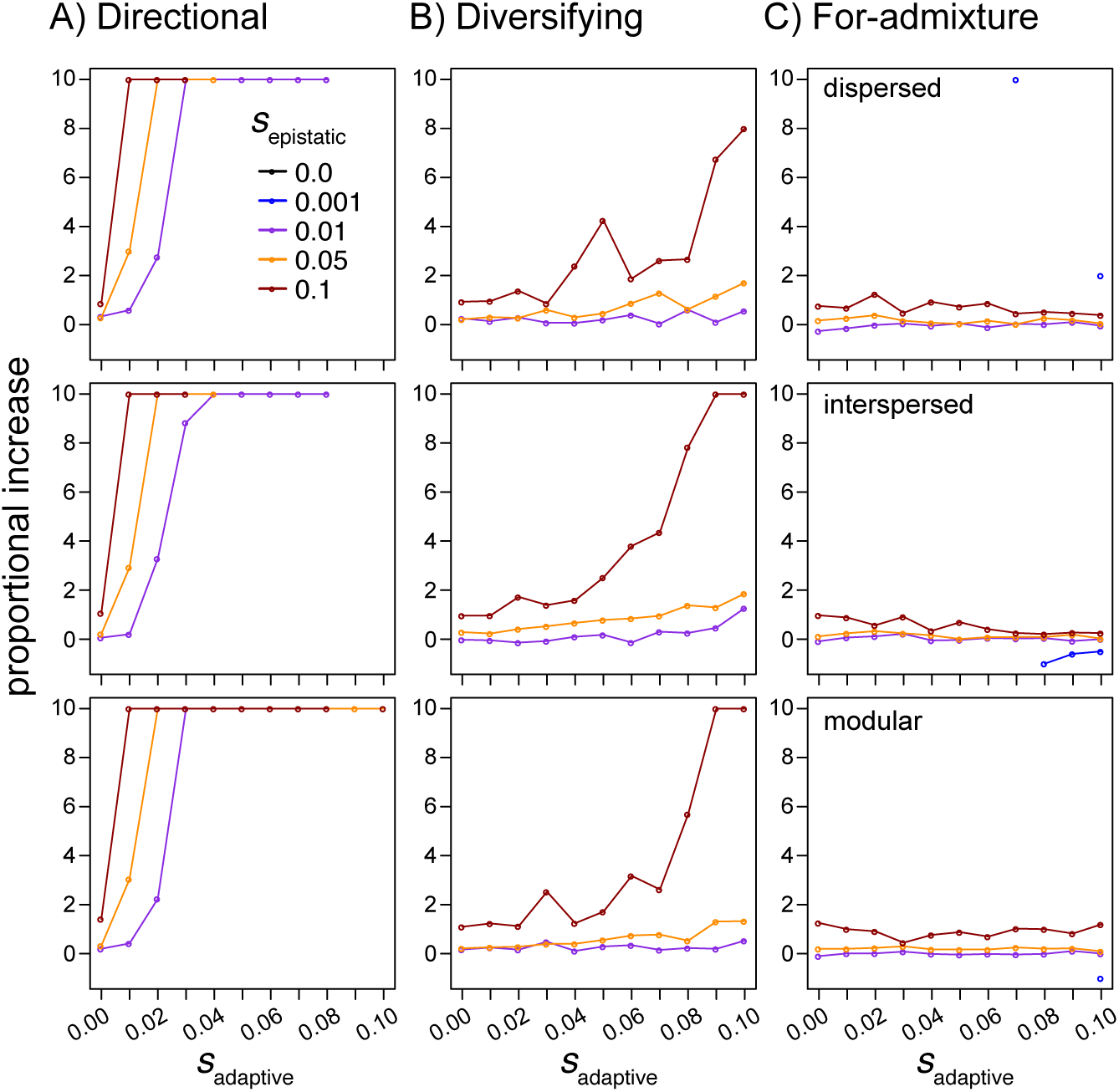
The effect of a bout of forced hybridization on the fraction of populations evolving RI. Proportional change (y-axis) was calculated as the difference in the number of populations evolving RI when simulations were initiated with F_1_S versus randomly mating parental individuals over the number of populations that evolved RI when simulations were initiated with randomly mating parental individuals. Symmetrical incompatibilities, in general, evolved more frequently in simulations initiated with a hybrid deme composed of F1 hybrid individuals compared to when initiated with equal proportions of randomly-mating parental genotypes. Results are shown for each of the three simulated genetic architectures (panel rows) under the directional selection (A), diversifying selection (B) and selection-for-admixture (C) models of selection acting on ‘adaptive’ loci. Recombination rates and migration were held at 0.2 and 0.001, respectively. In instances when there was a greater than 10-fold increase in the proportion of populations that evolved RI, values were rounded down to 10. Missing points occur for parameter combinations where no populations evolved RI across simulations initiated under either initial condition.

When populations are subject to an initial bout of hybridization, genetic architecture also has a larger effect on the probability of evolving RI. For example, a more modular architecture with weaker linkage between epistatic and adaptive loci is more permissive to symmetrical incompatibilities evolving under the directional and diversifying selection models (Figures S5 and S6, respectively). This is because a high frequency of F_1_ individuals helps to facilitate the formation of recombinant haplotypes, with the probability of a crossover events between different ‘types’ of loci being a function of their position along a chromosome. Modular architectures where loci are not in tight linkage are therefore the most conducive to the evolution of symmetrical incompatibilities when selection on adaptive loci is directional or diversifying (Figures S5 and S6), while linkage between adaptive and epistatic loci is more conducive to the evolution of symmetrical incompatibilities when selection favors admixture (Figures S7).

## Conclusions

Genome-wide sequence data has led to an increased appreciation of the prevalence of admixture and introgression between species (Payseur & Rieseberg 2016; Pease *et al*. 2016; Wallbank *et al*. 2016). While the consequences of hybridization have historically been viewed as maladaptive (Fisher 1930), others have proposed that hybridization can be a generative force that facilitates adaptive evolution and speciation (Seehausen 2004; Mallet 2007; Hedrick 2013; Nieto Feliner *et al*. 2017). If this is the case, hybridization may play a significant role in the production of biodiversity (Mallet 2007), and a few empirical examples have even linked the evolution of RI, without a change in ploidy, to hybridization and admixture occurring between different species (Rieseberg *et al*. 1995; Ungerer *et al*. 1998; Jiggins *et al*. 2008; Melo *et al*. 2009; Lamichhaney *et al*. 2017). Ascribing a causative role to hybridization and admixture in generating RI is however challenging, and the prevalence of HHS still remains largely unknown (Schumer *et al*. 2014).

Here I have focused on one general mechanism that can lead to the evolution of RI in hybrid populations: the fixation of different parental alleles at two or more groups of ‘coadapted’ or interacting loci (Buerkle *et al*. 2000; Schumer *et al*. 2015). Through simulation, I have shown that the evolution of RI due to symmetrical incompatibilities is strongly affected by (1) the strength and form of selection acting on different types of loci, (2) linkage relationships between adaptive and epistatic loci, (3) the arrangement of those loci along a chromosome, (4) gene flow between populations of hybrids and their parental species, and (5) the degree of hybridization occurring in a hybrid zone. These results suggest that there will be ‘sweet-spots’ – both genetic and ecological – that will be most conducive to the evolution of RI in hybrid populations. From a genetic perspective, weak incompatibilities between parental genomes are only capable of generating weak RI due to symmetrical incompatibilities. By contrast, strong and pervasive (in terms of number) incompatibilities will reduce the probability that admixed haplotypes will form and increase in frequency within a population. Therefore, the evolution of symmetrical incompatibilities will be most likely when parental species display an intermediate level of incompatibility; this will allow selection to maintain linkage disequilibrium between ‘coadapted’ alleles but not severely limit the ability of recombinant haplotypes to be present at an appreciable frequency within a population.

From an ecological perspective, the evolution of symmetrical incompatibilities is most likely when selection favors hybrid and admixed genotypes. Previous empirical work has shown that hybrid species tend to show novel ecologies or phenotypes when compared to their parental species (e.g. *Helianthus* sunflowers: (Rieseberg *et al*. 1995) *Heliconius* butterflies: (Melo *et al*. 2009; Salazar *et al*. 2010), *Geospiza* finches: (Lamichhaney *et al*. 2017)). These novel ecologies and phenotypes may be required to afford recombinant genotypes the opportunity to establish and evolve RI from their parental species, especially in a situation where hybrid populations are not found in geographic isolation.

Future work in speciation will benefit from continuing to quantify the extent of admixture within regions of hybridization and ultimately measure the fitness of hybrids relative to their parental species. Collecting these types of data across taxa that differ in the nature of hybridization (e.g. the extent of genetic divergence between parental species) and across a variety of environments are not trivial tasks. However, these data are needed if we are to understand the consequences of hybridization between species and populations in nature, and when and where we might expect to see admixed genomes stabilize and hybrid species evolve.

## Acknowledgements

I thank three anonymous reviewers, the subject editor at Molecular Ecology C. Alex Buerkle, and D. R. Matute, M. Servedio, and M. Schumer for helpful and constructive comments and/or discussions pertaining to previous versions of this manuscript. This work was supported by a NIH award R01GM121750 to D.R. Matute. I declare no conflict of interest with respect to any of the data or results herein presented.

## Data Accessibility

The scripts used to simulate hybrid populations are freely available at https://github.com/comeaultresearch/simuHybrid and will be deposited on Dryad upon acceptance. Files containing allele frequencies, recorded every 10 generations within the simulated populations analyzed here, will be deposited on Dryad upon acceptance.

## Author Contributions

AAC designed the study, analyzed the data, and wrote the manuscript.

